# Evidence for cPLA2 activation in Alzheimer’s Disease Synaptic Pathology

**DOI:** 10.1101/2025.03.27.645605

**Authors:** Qiu-Lan Ma, Brandon Ebright, Boyang Li, Jing Li, Jasmin Galvan, Ashley Sanchez, Michael Renteln, Dante Dikeman, Shaowei Wang, Bilal E. Kerman, Paul Seidler, Berenice Gutierrez-Grebenkova, Brooke E. Hjelm, Debra Hawes, Anne E. Hiniker, Kyle M. Hurth, David A. Bennett, Stan G. Louie, Helena C. Chui, Agenor Limon, Zoe Arvanitakis, Hussein N. Yassine

## Abstract

**Background:** Synapses are essential for learning and memory, and their loss predicts cognitive decline in Alzheimer’s disease (AD). Synaptic loss is associated with excitotoxicity, neuroinflammation, amyloid-β, and tau pathology, but the molecular mechanisms remain unclear. There is an urgent need to identify new targets to modify the disease and slow synaptic loss and cognitive decline. This study examines if calcium-dependent phospholipase A2 (cPLA2) is implicated in AD synaptic loss. cPLA2 catalyzes membrane phospholipids to release arachidonic acid, which can be metabolized into inflammatory eicosanoids.

**Methods:** cPLA2 levels were examined in synaptosomes isolated from the postmortem frontal cortex of individuals with no cognitive impairment (NCI), mild cognitive impairment (MCI), and AD dementia from the Religious Orders Study (ROS). Eicosanoids in synaptosomes were analyzed using lipidomics. Immunofluorescent staining investigated cPLA2 interactions with synaptic markers. Human iPSCs-derived neurons were used to study cPLA2 overactivation after exposure to amyloid-β 42 oligomers (Aβ42O), its relationships with synaptic markers, and the effects of cPLA2 inhibitors.

**Results:** We observed elevated cPLA2 (cPLA2α and cPLA2β) in AD synaptosomes and positive correlations with postsynaptic density protein 95 (PSD-95) and cognitive dysfunction. Eicosanoids were increased in AD synaptosomes and correlated with cPLA2, indicating cPLA2 activity at synapses/synaptosomes. Phosphorylated cPLA2α (p-cPLA2α) colocalized with PSD-95 in synaptosomes, and with postsynaptic Ca^2+^/calmodulin-dependent protein kinase IIα (CaMKIIα) and dendritic microtubule-associated protein 2 (MAP2) in NCI and AD brains, where their levels were reduced in AD. P-cPLA2α colocalizes with MAP2 at the neuronal soma associated with neuritic plaques and neurodegeneration in AD. Aβ42O activates cPLA2α in human iPSCs-derived neurons, leading to p-cPLA2α relocation from the cytosol to synaptic and dendritic sites to colocalize with CaMKIIα and MAP2, resulting in their reduction. P-cPLA2α also colocalized with PSD-95 in Aβ42O-exposed neurons, accompanied with increased PSD-95 intensity at soma membrane. These processes were reversed by the cPLA2 inhibitor ASB14780.

**Conclusions:** cPLA2 overactivation at synapses, dendrites, and excitatory neuronal somas is associated with synaptic loss, neuritic plaques and neurodegeneration, potentially contributing to cognitive decline in AD. Future research needs to explore the role of cPLA2 as a disease-modifying target for AD.

## Background

Synapses, which contain various neurotransmitters and neuropeptides, are essential for neural communications and play a vital role in learning and memory^1^. In Alzheimer’s disease (AD), synaptic loss serves as an early sign of memory impairment and cognitive decline^2^. This synaptic loss has been linked to multiple pathological conditions in AD, including excitotoxicity, oxidative stress, neuroinflammation, energy deficiency caused by mitochondrial dysfunction, amyloid plaque formation, and the development of neurofibrillary tangles. Despite these known associations, the disease-modifying targets that mediate synaptic loss in AD remain unclear. The current AD treatments, like donepezil and memantine, act at synaptic sites but only alleviate symptoms without affecting disease progression. There is an urgent need to identify new targets to modify the disease and slow synaptic loss and cognitive decline. This study presents evidence that a key phospholipid enzyme, calcium-dependent phospholipase A2 (cPLA2), is a potential disease-modifying target for AD.

cPLA2 hydrolyzes membrane phospholipids to release free arachidonic acid (AA) and lysophospholipids^3–5^. AA can be further metabolized by cyclooxygenases or lipoxygenases into proinflammatory prostanoids^6,7^. Among cPLA2 isozymes, cPLA2α (Group IVA) is the most studied enzyme due to its implication in inflammation in various diseases^8–12^. cPLA2α activation occurs through both enzymatic and non-enzymatic pathways. Several kinases can induce cPLA2α phosphorylation at multiple sites, including Ser505. These cPLA2α kinases include mitogen-activated protein (MAP) kinases^13^, Cdk5/p25^14,8,15^ and calcium/calmodulin-dependent protein kinase II (CaMKII)^16^. Notably, these kinases also phosphorylate tau^17–19^, a key pathological hallmark of AD. The cPLA2 enzyme contains an N-terminal C2 domain that regulates calcium-dependent activation^20–22^. Other factors capable of activating cPLA2α include Aβ^23–25^, lipopolysaccharide^26^, and high fat diet/obesity^27,28^. Studies on AD animal models have indicated that inhibiting cPLA2α leads to reduced tau phosphorylation^29^, improved synaptic and memory function^30^, and decreased Aβ pathology^31^. However, the impact of cPLA2 on human AD brain has been understudied. Studies have primarily reported elevated cPLA2α in astrocytes in AD^11,12,32^, but the impact of cPLA2 on human synapses and cognitive function remain largely unknown. Translational studies involved in understanding cPLA2 action in human brain synapses/synaptosomes are needed to support the development and efficacy of new brain penetrant cPLA2 inhibitors^33^.

To investigate cPLA2’s impact on human synapses and cognitive function, we studied synaptosomes isolated from postmortem human brain tissues in the Religious Orders Study (ROS), a well-characterized, prospective clinical and pathological cohort study^34–36^. Synaptosomes (also named synaptoneurosomes) are subcellular fractions that are abundant in fundamental, low-concentration proteins, and contain the basic structures and molecular machinery required for neurotransmission. They remain metabolically active and preserve functions such as plasticity and ion current signaling amenable to electrophysiological characterization, thus serving as excellent *ex vivo* models for investigating human synapses^37–41^. Therefore, we examined cPLA2 and eicosanoids levels in synaptosomes and analyzed its relationships with key synaptic protein markers involved in regulating memory processes, and cognitive performance in a group of deceased individuals from the ROS cohort, matched on clinical diagnosis. In parallel, we used human induced pluripotent stem cells (iPSCs)-derived neurons to study cPLA2 abnormal activation after exposure to amyloid-β 42 oligomers (Aβ42O), its relationships with synaptic markers, and the effects of cPLA2 inhibitors.

## Materials and methods

### Chemicals and reagents

All chemicals used in this study were purchased from Millipore Sigma, unless otherwise stated. Trans-Blot Turbo RTA Mini 0.2 µm Nitrocellulose Transfer Kit was purchased from Bio-Rad (#1704270). Laminin (23017015), Micro BCA™ Protein Assay Kit (#23235), and CyQUANT™ LDH Cytotoxicity Assay kit (#C20300) were purchased from ThermoFisher Scientific. Matrigel was purchased from Corning (# 354230). STEMdiff™ SMADi Neural Induction Kit (# 08581), STEMdiff™ Neural Progenitor Medium (#05833), and mTeSR™ Plus (# 100-0276) were purchased from StemCell Technologies. Antibodies were used in Table S3 antibody list.

### Human postmortem brain tissues and sections and clinical diagnoses

The use of human postmortem brain tissues and sections in this study were approved by an Institutional Review Board (IRB) of Rush University Medical Center and the University of Southern California. Participants in the ROS enrolled without known dementia and agreed to annual detailed clinical evaluation and brain donation. All participants signed informed and repository consents and an Anatomic Gift Act. The dorsolateral and medial prefrontal cortex (Brodmann area 9) of postmortem brain tissues from sixty individuals were obtained, which included individuals with no-cognitive impairment (NCI, n=20), mild cognitive impairment (MCI, n=15) and AD (n=25) as previously reported^34–36^. These samples had an average postmortem interval (PMI) time less than 5 hours which is within the 12 hours PMI period where synaptosomes isolated from human postmortem brain tissues, retain their metabolic activity, membrane potential, and are able to store, release, and reuptake neurotransmitters^41^. At death, the subjects underwent brain autopsy and postmortem assessment for AD pathology including neuropathologic criteria^34^. Detailed clinical and pathological data were also available (**Table S1**). Additional human postmortem brain sections were obtained from the University of Southern California (USC), Alzheimer’s disease research center neuropathology core, including subjects with NCI (n=4) and AD (n=3) (**Table S2**). The pathological assessments are performed based on the National Alzheimer’s Coordinating Center neuropathology form version 11.

### Isolation of synaptosomes from frozen postmortem brain tissues

Synaptosomes were isolated from frozen postmortem frontal cortices of ROS subjects, using a previously established protocol with minor modifications^32,42^. Briefly, brain specimens (∼100 mg) were homogenized in 10 volumes (w: v) of ice-cold homogenization buffer (10mM HEPES, pH 7.4, 0.32 M sucrose, 0.1 mM EDTA containing EDTA-free protease inhibitor cocktail (Roche, 04693159001) using a Teflon/glass homogenizer (10∼15 strokes). The homogenates were centrifuged at 1000 x g for 10 min to remove nuclear fractions, and the supernatants were centrifuged at 15,000 x g at 4°C for 30min to pellet the synaptosomes (P2 fraction). The synaptosomes were washed twice at 4°C in 1mL of ice-cold oxygenated Kreb’s-Ringer (K-R) solution with EDTA-free protease inhibitor cocktail. The synaptosomes were then resuspended in 400 μl of K-R solution, and the protein concentrations were determined by the Micro BCA™ Protein Assay.

### Isolation of gliosomes from frozen postmortem brain tissues

Gliosomes were isolated from frozen postmortem frontal cortices of subjects in the ROS cohort during synaptosome isolation process. Briefly, brain specimens (∼100 mg) were homogenized in 10 volumes (w: v) of ice-cold homogenization buffer (10mM HEPES, pH 7.4, 0.32 M sucrose, 0.1 mM EDTA containing EDTA-free protease inhibitor cocktail (Roche, 04693159001) using a Teflon/glass homogenizer (10∼15 strokes). The homogenates were centrifuged at 1000 x g for 10 min to remove nuclear fractions, and the supernatants were centrifuged at 15,000 x g at 4°C for 30min, the supernatants were collected as gliosomes (P1 fraction). The protein concentrations were determined by the Micro BCA™ Protein Assay.

### Lipid extraction and LC-MS quantification

The extraction, detection, and analysis of lipids from synaptosomes isolated from frozen postmortem brain tissues of ROS by LC-MS/MS lipidomics assay were performed as previously^43^. 30 μg of synaptosomes were used for the assay.

#### Single-Cell RNA Sequencing

Processed single-nucleus transcriptomic data and ROSMAP metadata were downloaded from the Synapse AD Knowledge Portal (https://www.synapse.org/#!Synapse:syn52293417) with Synapse ID: syn52293433 (Fig. 7g). Source data were collected from 427 (syn52293433) subjects from ROSMAP. Nuclei were isolated from frozen postmortem brain tissues and subjected to droplet-based single-nucleus RNA sequencing (snRNA-seq). Cell types were assigned based on Leiden clustering, marker gene analysis, and comparisons with previously published data in the original publication.

#### Gene Set Enrichment Analysis

The R package GSVA (method option “ssGSEA”) was used to calculate the pathway enrichment score of each subject. Single sample GSEA (ssGSEA) is a non-parametric method that calculates the normalized difference in empirical cumulative distribution functions (CDFs) of gene expression ranks inside and outside the gene set, representing the enrichment score for the gene set. The annotated gene sets were retrieved from MsigDB. REACTOME_ARACHIDONIC_ACID_METABOLISM (M27140) was selected to represent AA metabolism. REACTOME_ARACHIDONIC_ACID_METABOLISM GENE_SYMBOLS PON1,ALOX5,DPEP1,TBXAS1,PTGS2,PTGS1,GGT5,GGT1,ABC C1,PON3,PON2,PTGR1,PTGDS,ALOX12,PTGES3,LTA4H,PLA2G4A,FAAH,EPHX2,P TGIS,ALOX5AP,CYP2J2,CYP1B1,CYP2C9,CYP2C8,PTGR2,CYP1A1,CYP1A2,DPEP3 ,CYP4B1,PTGES2,PTGES,CYP2U1,PRXL2B,CBR1,ALOX15,CYP4A22,MAPKAPK2,H PGDS,HPGD,FAAH2,CYP2C19,DPEP2,GPX4,CYP4F11,CYP4F22,GPX2,ALOXE3,AL OX12B,ALOX15B,CYP8B1,CYP4F2,CYP4F8,CYP4F3,CYP4A11,AKR1C3,AWAT1,LTC 4S,GPX1,ALOX5,ABCC1,LTC4S

### Transmission electron microscope (TEM)

To validate the morphological integrity of the isolated synaptosomes, 6 μl of isolated synaptosomes were mixed with 6 μl 4% Uranyl acetate solution, then the images were taken under TEM.

### Lactate dehydrogenase activity (LDH) assay in synaptosomes

To evaluate synaptosomes membrane integrity, LDH activity test was conducted as previously^32,44^. Briefly, the isolated synaptosomes were incubated in 1% Triton X-100 lysis buffer for 1 hour on ice. Then, 50 μl of synaptosome in KR buffer with 1% Triton X-100 (total LDH) or without Triton X-100 (free LDH) were incubated with 50 μl LDH reaction mixture for 30 minutes. After adding the stop solution, the absorbance at 490 nm and 680 nm were measured. The LDH activity was indicated as the absorbance at 490–680 nm.

### Immunofluorescent staining synaptosomes

15 μg of synaptosomes were used for immunofluorescent staining of the synaptic marker PSD-95 and phospho-cPLA2α (p-cPLA2α). Briefly, the extracted synaptosomes were washed with 200 µl of TBS by gentle pipetting and then centrifuged at 15,000 x g for 10 mins at 4°C. The resulting synaptosome pellet was fixed with 200 µl of ice-cold 95% ethanol for 5 mins at 4°C. After fixation, the synaptosomes were briefly washed with TBS and incubated with blocking buffer (5% BSA, 5% goat serum, and 0.3% Triton-X in TBS) for 1 hour at room temperature. Following blocking, the synaptosomes were incubated with the primary antibodies at 4°C overnight. After three washes with TBS, the samples were incubated with secondary antibodies at room temperature for one hour. Images were acquired using a LSM 800 Zeiss and SP8 Leica confocal microscope.

### Western blotting

For Western blotting (WB) detection of assessed antigens, synaptosome protein concentration was determined using the BCA assay. Equal amounts of protein per sample were added to Laemmli loading buffer, and boiled for 5 min. 5∼15 µg of protein per well was electrophoresed on 10% Tris-glycine gels and transferred to nitrocellulose membranes. The WB following processes were same as previously^32^.

### Immunofluorescent staining and image analysis

The immunofluorescent staining methods were used as previously described^45^.Formalin-fixed and paraffin-embedded postmortem brain sections from USC ADRC neuropathological Core were used to examine p-cPLA2, MAP2, and CamKIIα in human brains. All images were taken with a LSM 800 Zeiss and SP8 Leica confocal microscope and processed using CellProfiler or ImageJ.

### Generation of human induced pluripotent stem cells (iPSCs)-derived neurons and treatment by amyloid-beta (Aβ)42 oligomers with or without ASB14780

Neural progenitor cells (NPCs) were differentiated from iPSCs using Neural Induction Kit following manufacturer’s protocol. Briefly, the JAX IPSC 1150 (APOE4/4) iPSC line was grown on Matrigel-coated dishes in mTeSR™ Plus. To initiate NPC induction, iPSCs were passaged to either Matrigel- or Poly-L-Ornithine hydrobromide/laminin-coated plates and were fed with Neural Induction Medium + SMADi. At passage 3, cells were switched to STEMdiff™ Neural Progenitor Medium and were maintained in that medium. For neuronal differentiation, NPCs were seeded on PORN/laminin-coated chamber slides at 20,000 cells/cm^2^. For neuronal induction, cells were fed with the neuronal medium as described in Bardy et al^46^. At five weeks of neuronal differentiation, cells were treated with 2.5 μM Aβ42O with or without cPLA2 inhibitor ASB4780 for 72 hours, then the cells were fixed in 4% PFA for 15 mins and stained for p-cPLA2α, PSD-95, MAP2, and CaMKIIα.

### Statistical analysis

Statistical analyses were performed with PRISM 6.0 software. Differences among the means ± SEM were assessed by ANOVA followed by Tukey-Kramer post hoc tests. Statistical significance was *a priori* defined as *p* < 0.05.

## Results

### Decreased synaptosome yield in individuals with MCI and AD (ROS)

We observed a significant decrease in synaptosome yield extracted from the frontal cortex of individuals with mild cognitive impairment (MCI, n=15) and AD (n=25) compared to those with no cognitive impairment (NCI, n=20) **(\****p* < 0.05, **Fig. 1a).** No differences in the yield of gliosomes and myelin were observed between the groups (*p* > 0.05, **Fig. 1b, c**). These results indicate a loss of synapses/synaptosomes but not myelin in patients with MCI and AD. Clinical and pathological characteristics of the ROS samples are shown in **Table S1**^47,48^.

**Fig. 1.**
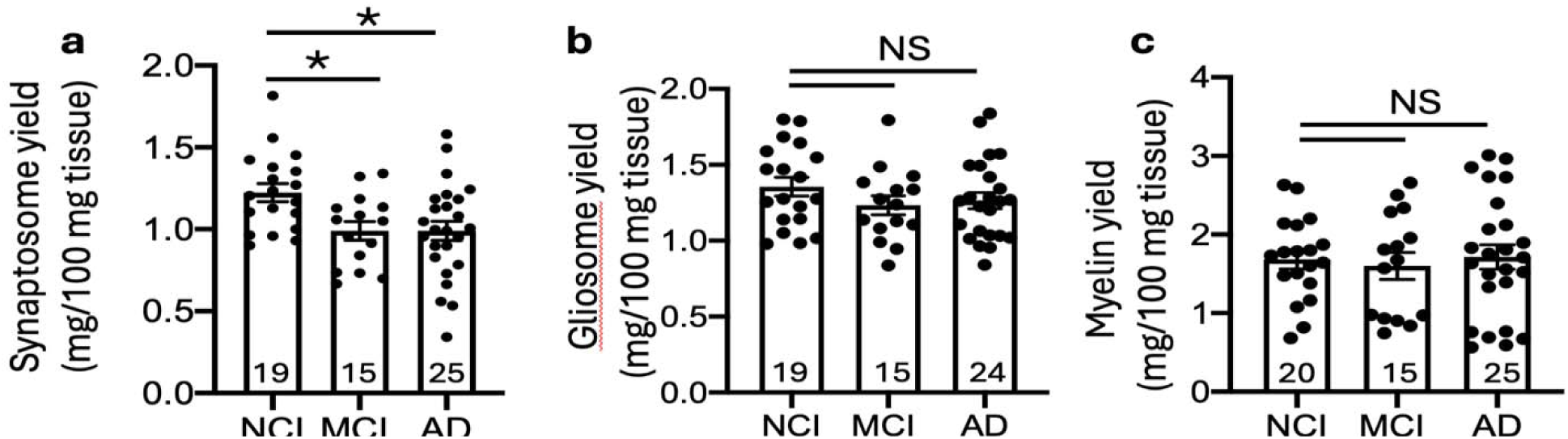
Reduction of synaptosome yield in MCI and AD cases in the ROS. **a.** Compared to NCI, synaptosome yield was reduced in MCI and AD. One-way ANOVA, Tukey’s multiple comparisons test, *F*_(2, 56)_ = 5.250, *P* = 0.0081. **b.** There was no differences for gliosome yield among groups. One-way ANOVA, Tukey’s multiple comparisons test, *F*_(2, 55)_ = 1.076, *P* = 0.3479. **c.** There was no differences for myelin yield among groups. One-way ANOVA, Tukey’s multiple comparisons test, *F*_(2, 56)_ = 0.1334, *P* = 0.8754. Data are presented as mean ± SEM, \**P* < 0.05.

The specificity and integrity of the isolated synaptosomes from frozen postmortem human brain tissue were validated by WB, LDH assay, IF staining, and transmission electron microscopy (TEM). WB analysis revealed higher levels of anti-PSD-95 antibody immunoreactivity in the synaptosome-enriched P2 fraction, but not in gliosome-enriched P1 fraction (**Fig. S1a)**. In contrast, anti-astrocyte marker GFAP and microglia marker IBA1 antibodies showed higher intensities in the P1 fraction. Synaptosomal integrity was confirmed by LDH assay (**Fig. S1b**), immunofluorescent staining using anti-PSD-95 antibody (**Fig. S1c**), and TEM (**Fig. S1d**). TEM also showed the loss of synaptic vesicles (SVs) within synaptosomes (Syn) in an 80-year-old female with AD (PMI 3.8 hours) compared to a 95-year-NCI female (PMI 3.2 hours, **Fig. S1d**).

### Increased cPLA2 in synaptosomes of MCI and AD (ROS)

The role of cPLA2α in astrocytes has been previously studied in AD, but its role in neurons of the human AD brain^11,12,32^ has not been explored. cPLA2β has not been studied in AD. Thus, we examined cPLA2 (cPLA2α and cPLA2β) in the synaptosomes. Compared to NCI, both cPLA2α and cPLA2β were increased in AD (\*\**p* < 0.01, **Fig. 2a, b)**. cPLA2β was also increased in MCI (\**p* = 0.0274, **Fig. 2b)**. A significant correlation between cPLA2α and cPLA2β was observed (r = 0.5268, \*\*\*\**p* < 0.0001, **Fig. 2c**), suggesting they may share functional roles as previously reported^49^. Phospho-cPLA2α (p-cPLA2α) showed a trend towards increased levels in AD (*p* = 0.1, data not shown). This WB analysis included 60 samples, however, two outliers were identified (denoted by arrow) and were excluded from the statistical tests. One outlier belonged to the MCI group and displayed the highest level of p-cPLA2α levels among samples; the other outlier, which belonged to the AD group, showed protein degradation. Specific bands for p-cPLA2α and cPLA2α were validated using cPLA2α knockout (KO) mice. Specific bands for cPLA2β was first detected by an anti-cPLA2 isoform antibody and further validated by an anti-cPLA2β specific antibody (**Fig. 2d**).

**Fig. 2.**
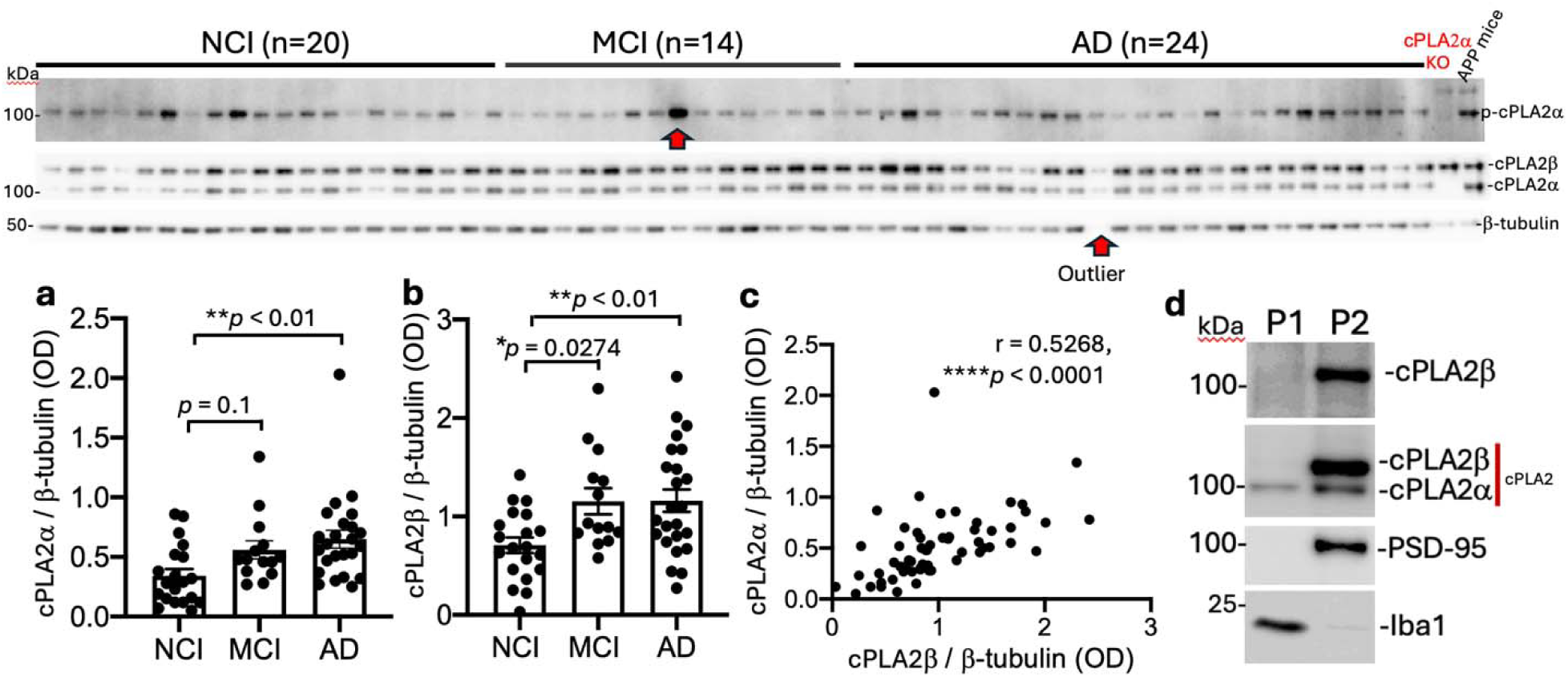
Elevation of cPLA2 in synaptosomes of AD in ROS. **a.** Compared to NCI, cPLA2α showed an increase in AD. One-way ANOVA, Tukey’s multiple comparisons test, *F*_(2, 55)_ = 5.477, *P* = 0.0068. **b.** cPLA2β showed an increase in both MCI and AD. One-way ANOVA, Tukey’s multiple comparisons test, *F*_(2, 55)_ = 5.698, *P* = 0.0056. **c**. cPLA2ß corelated with cPLA2α. The graph shows Pearson correlation between two variables, as described on the axis, \*\*\*\**P* < 0.0001. **d**. The enrichment of cPLA2β in synaptosomes (P2) was validated by a cPLA2β specific antibody. Data are presented as mean ± SEM., \**P* < 0.05, \*\**P* < 0.01, \*\*\*\**P* < 0.0001.

### Correlation of synaptic cPLA2 with measures of cognitive function

Next, we investigated the correlation between synaptic cPLA2 levels and clinical measures of cognitive function. Interestingly, **Fig.3a-g** shows a weak but significant inverse correlation between cPLA2 and different memory subdomains. p-cPLA2α was negatively correlated with global cognition (r = -0.2753, \**p* = 0.0365, **Fig.3a**) and episodic memory (r = -0.2724, \**p* = 0.0386, **Fig.3b**), suggesting that higher p-cPLA2α levels are associated with lower cognitive performance levels. cPLA2α was negatively correlated with working memory (r = -0.2919, \**p* = 0.0262, **Fig.3c**) and perceptual speed (r = -0.3491, \*\**p* < 0.01, **Fig.3d)**, suggesting that increased cPLA2α is linked to declines in these cognitive functions. cPLA2β was also negatively correlated with global cognition (r = -0.3662, \*\**p* < 0.01, **Fig.3e**), episodic memory (r = -0.3752, \*\**p* < 0.01, **Fig.3f**), and working memory (r = -0.3418, \**p* < 0.01, **Fig.3g**), further supporting the association between elevated cPLA2β levels and low cognitive performance. The correlation between cPLA2 and other memory domains was also analyzed, including semantic memory and visuospatial ability/perceptual orientation; however, no significant associations were identified (data not shown).

**Fig. 3:**
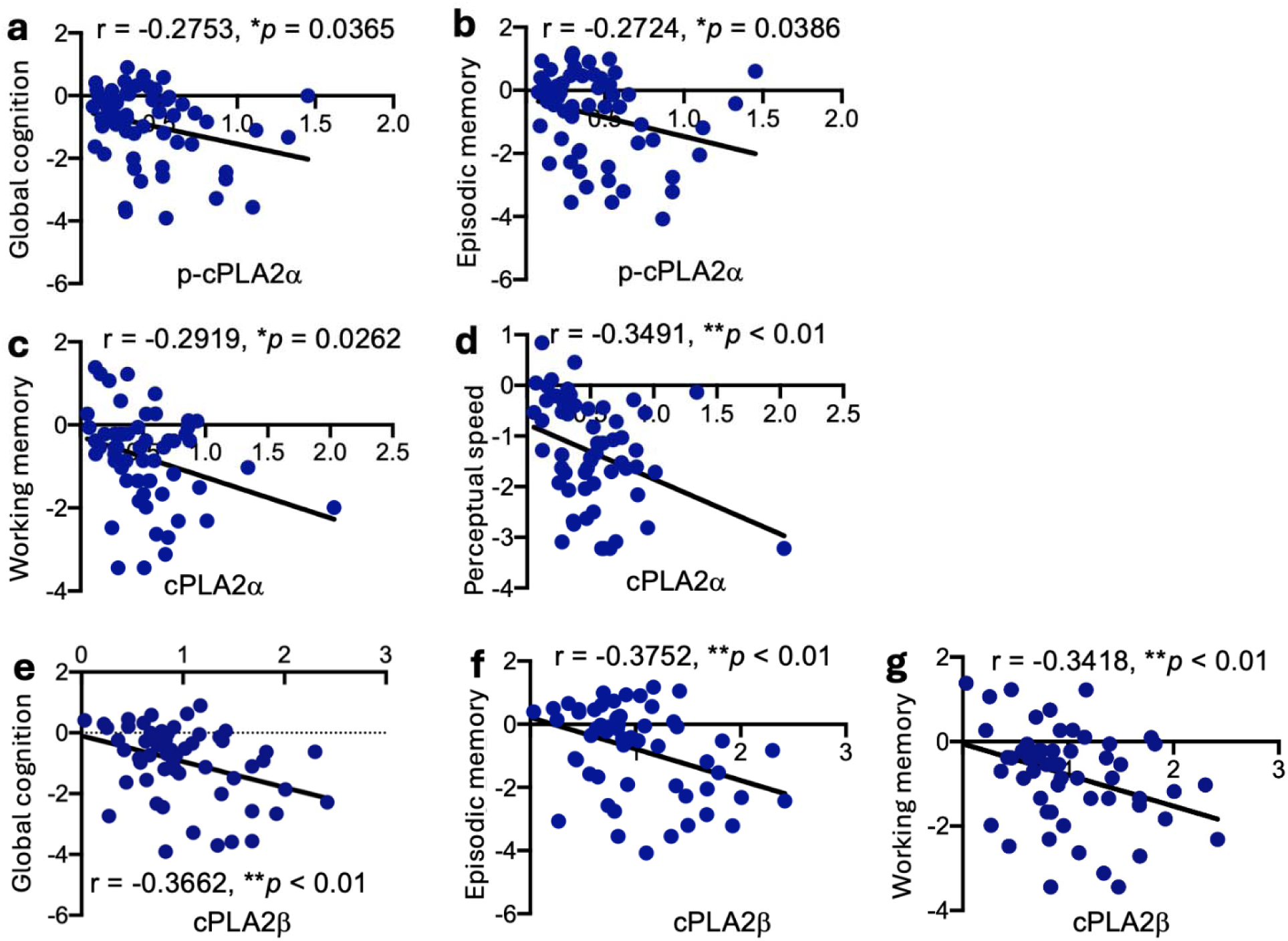
Correlation between cPLA2 in synaptosomes and cognitive dysfunction in ROS. **a**, **b.** p-cPLA2α was negatively correlated with global cognition, and episodic memory. **c**, **d.** Total cPLA2α levels was negatively correlated with working memory and perceptual speed. **e**-**g.** cPLA2β was negatively correlated with global cognition, episodic memory, and working memory. **a-g**. Graphs show Pearson correlation between two variables, as described on the axis, \**P* < 0.05, \*\**P* < 0.01.

### Correlation and interaction between PSD-95 and cPLA2 in synaptosomes

Postsynaptic density-95 (PSD-95) plays vital roles in learning and memory processes^50,51^. PSD-95 in synaptosomes was analyzed in the ROS participants using WB. **Fig. 4**, PSD-95 levels showed a trend increase in AD compared to NCI (p = 0.0988, **Fig.4a**), suggesting enhanced postsynaptic protein PSD-95 presence in AD synapses. PSD-95 levels were strongly correlated with cPLA2α (r = 0.6574, \*\*\*\**p* < 0.0001, **Fig.4b**), and cPLA2β (r = 0.7889, \*\*\*\**p* < 0.0001, **Fig.4c**), indicating a potential link between increased cPLA2 isoforms and PSD-95 protein density in synaptosomes. PSD-95 levels were also correlated with p-cPLA2α (r = 0.341, \*\**p* < 0.01, data not shown). b. Immunofluorescent staining (IF) revealed a colocalization between PSD-95 and p-cPLA2α (**Fig.4d**) in synaptosomes, demonstrating after the cytosolic cPLA2α was phosphorylated, it relocated to postsynaptic sites to interact with PSD-95.

**Fig. 4.**
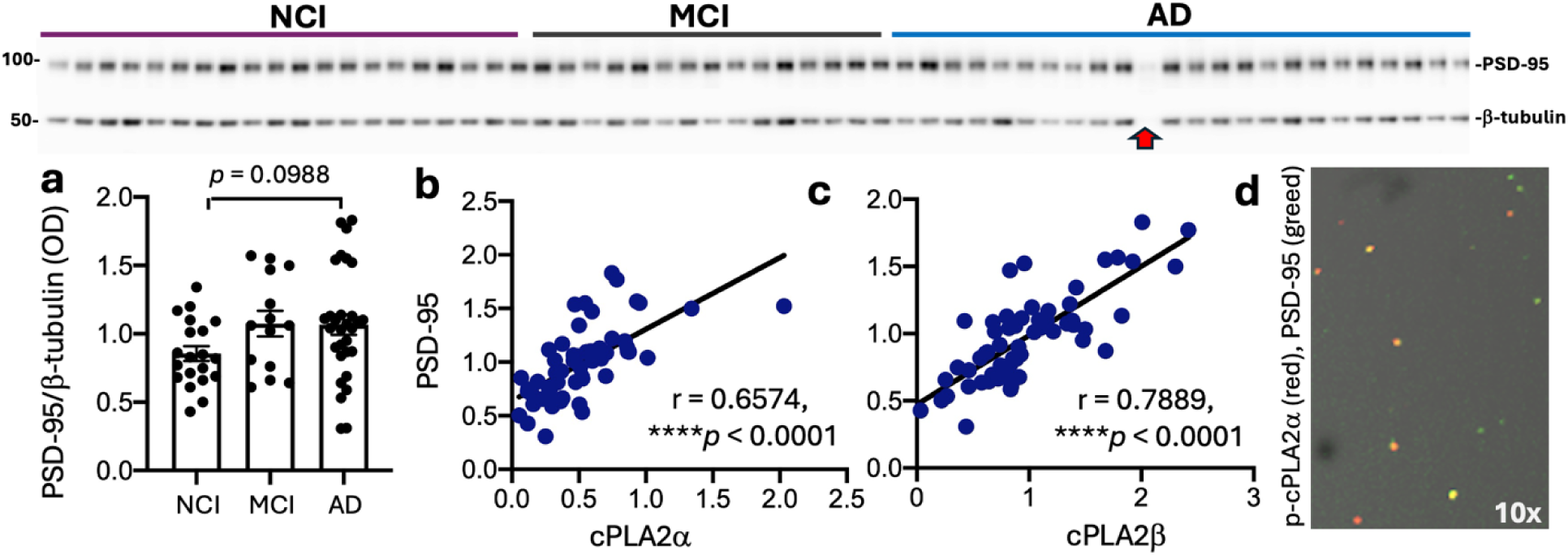
Correlation and interaction between cPLA2 and PSD-95 in synaptosomes (Religious Orders Study). **a.** PSD-95 shows a trend toward increased levels of synaptosomes from AD cases compared to NCI controls. One-way ANOVA, Tukey’s multiple comparisons test, *F*_(2, 61)_ = 2.583, *P* = 0.0838. Data are presented as mean ± SEM. **b, c.** PSD-95 levels are positively correlated with cPLA2α and cPLA2β. Graphs show Pearson correlation between two variables, as described on the axis, \*\*\*\**P* < 0.0001. **d.** IF staining shows a colocalization between p-cPLA2α and PSD-95 in synaptosomes.

### Reduction of CaMKIIα in synaptosomes of AD and its colocalization with P-cPLA2α in aging and AD brains

Calcium–calmodulin (CaM)-dependent protein kinase IIα (CaMKIIα) regulates PSD-95 activity^52^. It is the most abundant protein in excitatory synapses and is central to synaptic plasticity, learning and memory^53^. We examined CaMKIIα in the synaptosomes of participants in the ROS. **Fig 5 a, b.** WB analysis shows that p-CaMKIIα was increased in synaptosomes of MCI compared to NCI (*p* = 0.030), with a decrease in AD compared to MCI *(p* = 0.053), indicating dynamic changes in CaMKIIα phosphorylation during disease progression. Total CaMKIIα displays a trend toward increased levels in MCI compared to NCI, followed by a trend decrease in AD compared to MCI (*p* = 0.1, data not shown), suggesting alterations in synaptic CaMKIIα content in AD. P-CaMKIIα had a weak but significant correlation with cPLA2β (r = 0.281, \**p* = 0.034, **Fig 5c**). A colocalization of p-cPLA2α with CaMKIIα was observed (yellow color, **Fig. 5d)**. Compared to NCI, p-cPLA2α intensity was increased in AD, which was accompanied with a decrease in CaMKIIα (\*\*\*\**p*<0.0001, **Fig. 5e, f)**. These data indicated that activated cPLA2α interacts with CaMKIIα at postsynaptic membrane phospholipids, which potentially disrupts CaMKIIα signaling pathway.

**Fig. 5.**
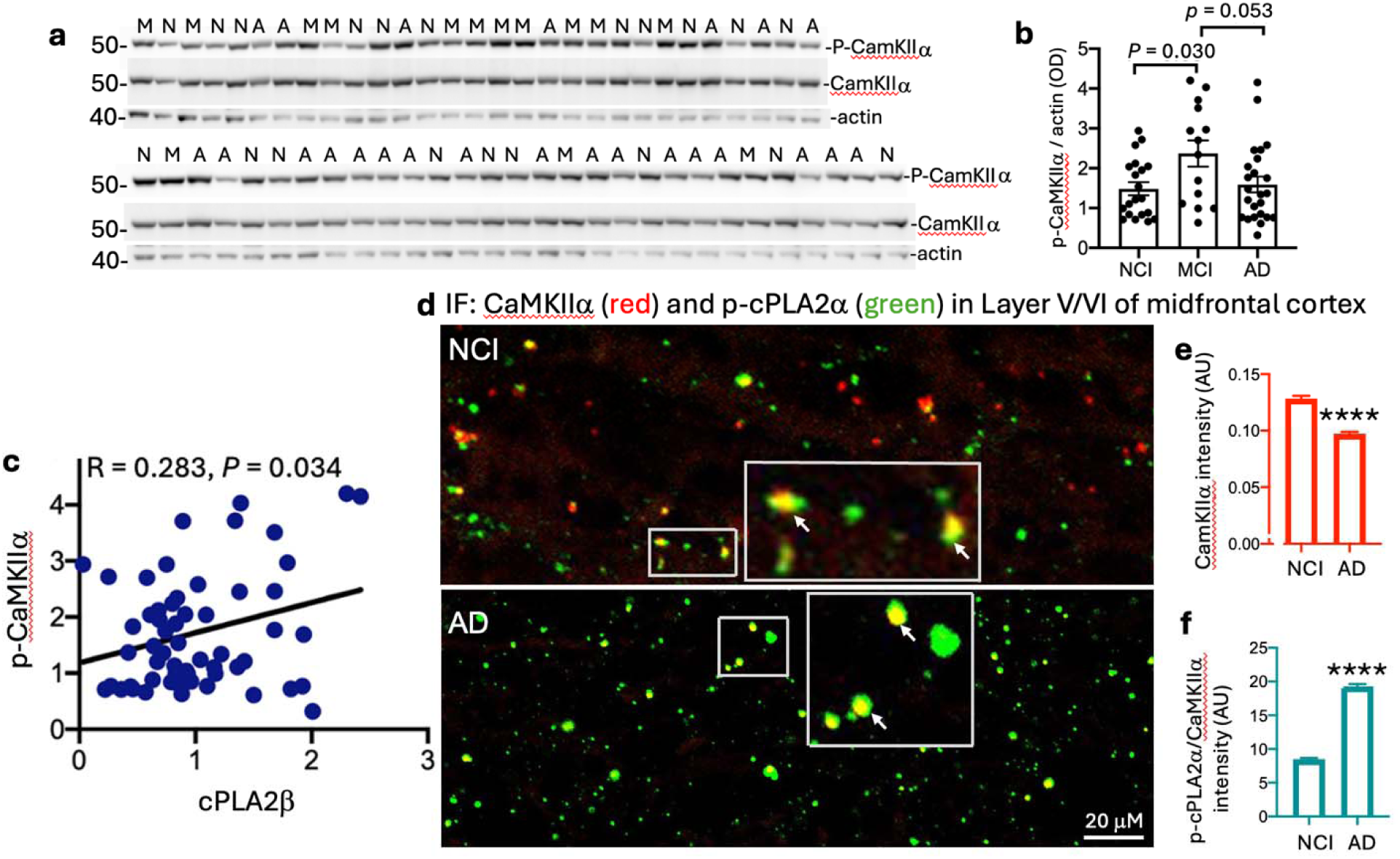
**Reduction of CaMKII**α **in synaptosomes of AD and its colocalization with P-cPLA2**α **in aging and AD brains. a, b.** P-CaMKIIα shows an increase in MCI compared to NCI, with a trend toward a decrease in AD compared to MCI. One-way ANOVA, Tukey’s multiple comparisons test. **c.** cPLA2β shows a weak but significant positive correlation with p-CaMKIIα. The graph shows Pearson correlation between two variables, as described on the axis, \**P* < 0.05. **d.** IF staining of p-cPLA2α (green) with CaMKIIα (red) showing their colocalization (yellow arrows) in layer V/VI of middle frontal cortex of AD brains. **e.** CaMKIIα intensity is decreased. **f.** P-cPLA2α intensity increased in AD, two-tailed unpaired Student’s *t-*test. We analyzed 2500 synaptic objects in NCI and 4515 in AD, identified using CellProfiler. Data are presented as mean ± SEM, \**P* < 0.05, \*\*\*\**P* < 0.0001.

### Reduction of dendritic MAP2 at synaptic sites and its colocalization with p-cPLA2α at neuronal soma membrane associated with neuritic plaques in AD brains

To further investigate the interaction between cPLA2 and dendritic/postsynaptic proteins, we performed double immunofluorescent staining of p-cPLA2α with MAP2 in both NCI and AD. P-cPLA2α was also colocalized with MAP2 in both NCI and AD (**Fig. 6a, d)**. However, when compared to NCI, the intensity of MAP2 at synaptic sites was reduced in AD (**Fig. 6b**), which was also accompanied by an increase in P-cPLA2α intensity (**Fig. 6c**). Additionally, the colocalization of P-cPLA2α with MAP2 at neuronal soma membrane is associated with core neritic plaques and degenerated neurons (MAP2 aggregation) in AD brain (**Fig. 6d)**. Taken together, these data suggest that in AD, activated-cPLA2α translocates to the postsynaptic/dendritic and neuronal soma membrane phospholipid sites, where it interacts with MAP2, promoting synaptic loss, neuritic plaques, and neurodegeneration.

**Fig. 6.**
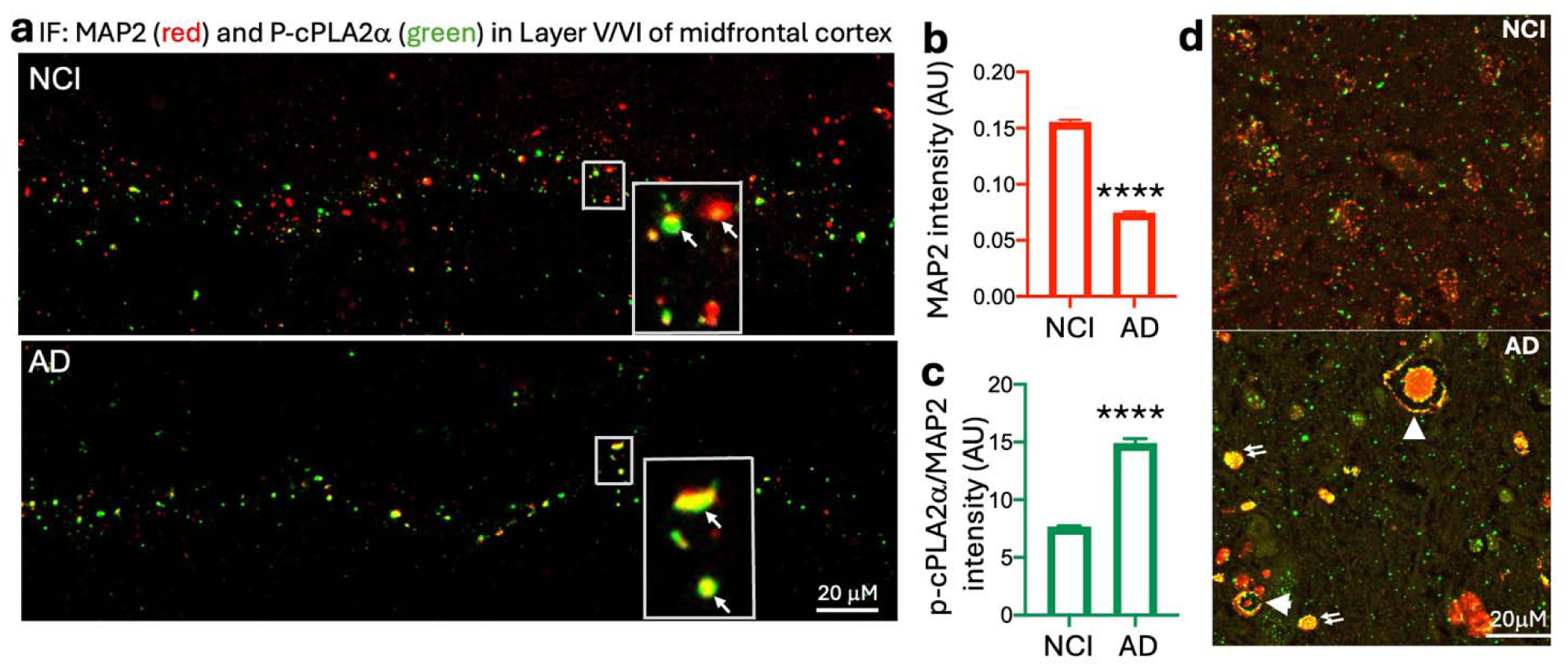
**Reduction of MAP2 at synaptic sites and its aggregation at neuronal soma is associated and colocalized with p-cPLA2**α **in AD. a-c.** IF staining shows p-cPLA2α (green) and MAP2 (red) colocalization (yellow arrows) in layer V/VI of middle frontal cortex in AD brains. P-cPLA2α intensity increased in AD but MAP2 intensity was decreased. \*\*\*\**P* < 0.0001. mean ± SEM, two-tailed unpaired Student’s *t-*test. We analyzed 9320 synaptic objects in NCI and 6465 in AD, identified using CellProfiler. **d.** Colocalizations of p-cPLA2a with MAP2 at neuronal soma membrane showed core neuritic plaques (double arrow) and degenerated neurons (arrow) in AD brain.

### Lipidomic analysis of AA and AA metabolites in synaptosomes and their correlation with cPLA2 and AD pathology

To evaluate cPLA2 activity in synaptic membrane phospholipids, we conducted a lipidomic profiling assay in synaptosomes of a subgroup with NCI and AD (n=11/group) from the ROS. Overall, relative to NCI, AD cases showed high levels of n-6 polyunsaturated fatty acids (PUFAs), AA, and AA metabolites, as shown in the heatmap (**Fig. 7a, b**). cPLA2α was highly correlated with AA metabolites of TXB2 (r = 0.846, \*\*\*\**p* < 0.0001) and PGF2α (r = 0.744, \*\*\*\**p* < 0.0001, **Fig. 7c, d**). AA was correlated with both cPLA2α (r = 0.5261, \*\**p* < 0.01, **Fig. 7e**) and cPLA2β (r = 0.5203, \**p* = 0.01, **Fig. 7f**). cPLA2β was also significantly correlated with PGD2 (r = 0.496, \**p* = 0.0161), PGE2 (r = 0.4623, \**p* = 0.0264), and adrenic acid (r = 0.438, \**p* = 0.0365, data not shown). These data are consistent with the functional role of cPLA2α and cPLA2β for releasing free AA and that can be further metabolized to form eicosanoids in synaptosomes. Additionally, we further analyzed AA metabolism pathway scores (surrogate of cPLA2 activation) in layer V/VI of excitatory neurons in relation to AD pathology in a study of brain tissues from the ROS and the Memory and Aging Project (ROSMAP) (n=427), utilizing single-cell RNA sequencing. The heatmap (**Fig. 7g**) illustrates the correlation between AA metabolism scores and various AD variables (columns) in layer V/VI of excitatory neurons using a linear mixed effect model. The color scale, from blue (negative correlation) to red (positive correlation), indicates the strength and direction of these correlations. Cognition and AD pathology have opposite scales, with strong correlations between greater AA metabolism with lower cognitive function and greater AD pathology in excitatory neurons. Taken together, these data further indicate higher cPLA2 activation at AD synaptosomes/synapses, with potentially significant and detrimental implications in synaptic loss, cognitive dysfunctions, and overall AD pathology.

**Fig. 7.**
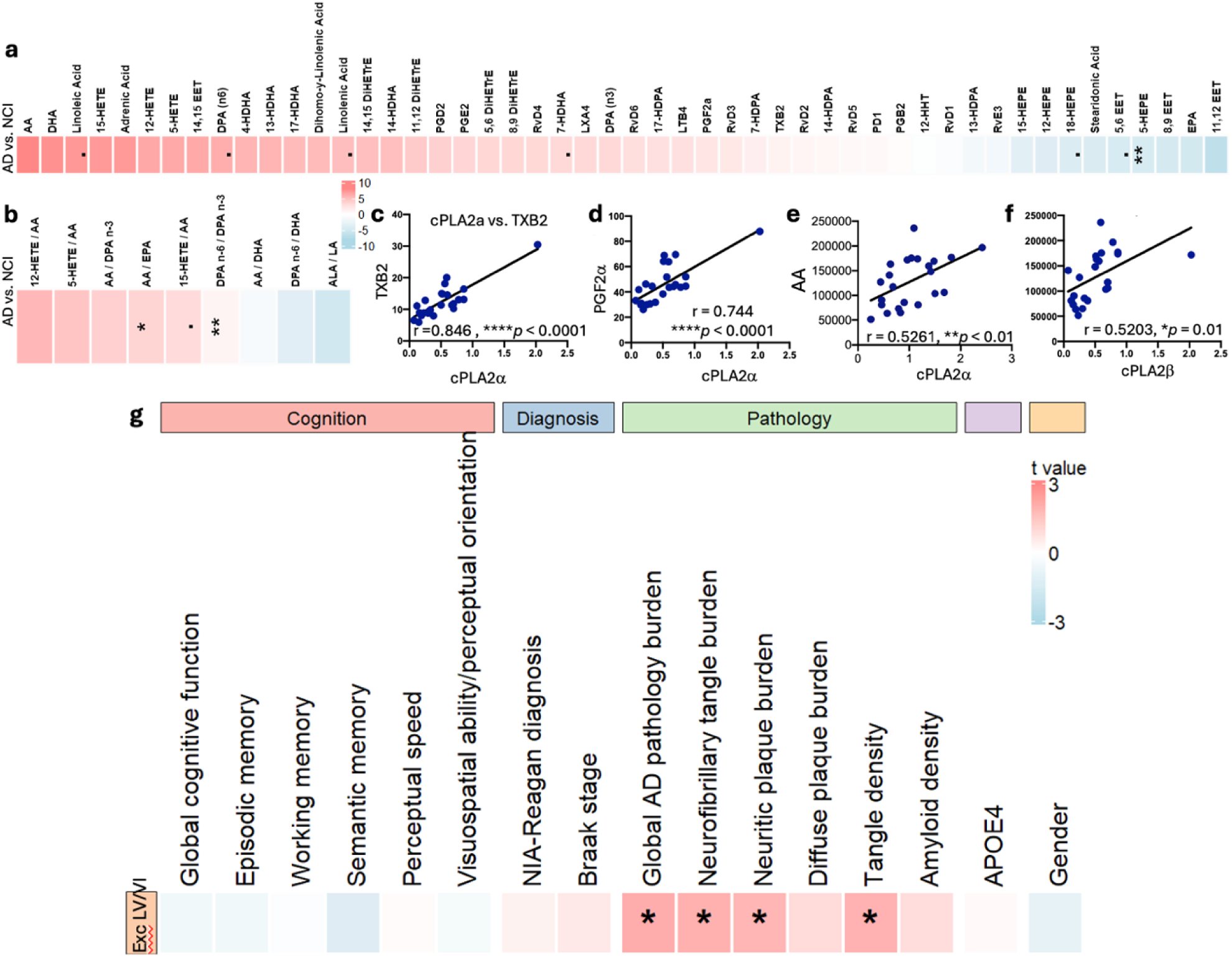
Increased arachidonic acid (AA) and AA metabolites in synaptosomes of AD and their correlation with cPLA2 and AD pathologies. a, **b.** Heatmap generated from lipidomic analysis of PUFAs and PUFAs’ metabolites in synaptosomes comparing AD vs. NCI by Wilcox rank sum test. The color represents the Wilcox test estimate, where redness indicated the increased lipid species in AD, and vice versa. Overall, relative to NCI, AD showed high arachidonic acid (AA) and AA metabolites, including increased TXB2 and PGF2α, which were highly corelated with cPLA2α (**c, d**). AA was corelated with both cPLA2α (**e**) and cPLA2β (**f**). **c-d** Graphs show Pearson correlation between two variables, as described on the axis, \**P* < 0.05, \*\**P* < 0.01, \*\*\*\**P* < 0.0001. **g.** SnRNA-seq analysis of 427 human brain tissues from ROSMAP shows strong correlations between AA metabolism scores in layer V/VI excitatory neurons with AD pathologies. The association is represented by the t-value (the effect size divided by standard error of effect size). Redness indicates significant positive associations, blueness negative associations. Significance is defined by an FDR (false discovery rate) adjusted p-value (*p < 0.1, *p < 0.05, **p < 0.01, ***p < 0.001*).

### Activated cPLA2α mediated Aβ42 oligomers-induced reduction of CaMKIIα in human iPSC-derived neurons, which was rescued by the cPLA2 inhibitor ASB14780

Since Aβ42O are one of the important factors driving synaptic loss in AD and AD models^54^, we examined the role of cPLA2α in Aβ42O-induced synaptic dysfunction in a mixed-culture of neurons and astrocytes derived from a human iPSC line carrying the APOE4/4 genotype. NPCs (neural progenitor cells) were generated using a commercially available kit. After NPCs were differentiated for five weeks^46^, the cells were exposed to 2.5 μM Aβ42O, with or without 1 μM of the cPLA2α inhibitor ASB14780 (ASB) for 72 hours. The chosen dosage of Aβ42O was determined through studies investigating the impact of soluble Aβ42 species on the synaptic dysfunction of neurons derived from human iPSCs^55^. Compared to untreated control cells, Aβ42O increased p-cPLA2αSer505 intensity at soma membrane (**Fig.8a, b**), suggesting the translocation of p-cPLA2a from the cytosol to membrane in neurons. In contrast, CaMKIIα intensity was reduced by Aβ42O treatment (**Fig.8a, c**), indicating the synaptic loss. These changes were partially reversed by treatment with cPLA2a inhibitor ASB14780. The colocalization between p-cPLA2a and CaMKIIα was observed in Aβ42O-treated neurons (arrow, yellow color). These processes were partially suppressed by cPLA2α inhibitor ASB14780. These data suggesting that cPLA2α meditates Aβ42O-induced synaptic loss and cPLA2α inhibitor ASB14780 partially protects the synapses from these processes.

**Fig. 8.**
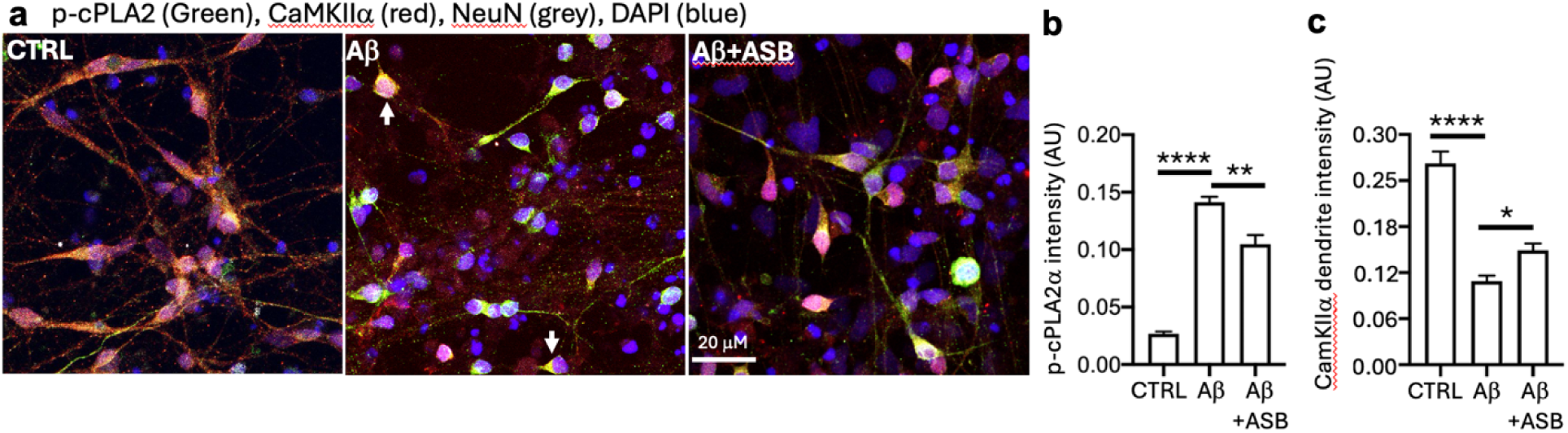
Aβ42 oligomers triggered cPLA2α activation and reduced CaMKIIα levels in dendrites, which was suppressed by the cPLA2 inhibitor ASB14780 in human iPSC-derived neurons. a.b. Compared to untreated control (CTRL) cells, Aβ42O (Aβ) reduced CaMKIIα-stained dendritic intensity in human iPSC-derived neurons. **a, c.** cPLA2α inhibitor ASB14780 (ASB) partially protected Aβ42O-induced reduction of CaMKIIα. The colocalization between CaMKIIα and p-cPLA2α was observed in Aβ42O-challenged neurons (yellow color, arrow, **a**). One-way ANOVA, Tukey’s multiple comparisons test. Data are presented as mean ± SEM, *\*p < 0.05, **p < 0.01, ***p < 0.001, ****p < 0.001*).

### cPLA2α activation mediated Aβ42 oligomer-induced increase in MAP2 and the colocalization in human iPSC-derived neurons, which was attenuated by ASB14780

To investigate if the activated cPLA2 affects MAP2 in Aβ42O (Aβ)-challenged human neurons derived from iPSCs, we further analyzed MAP2 from the same experiment. **Fig.9** shows that Aβ42O increased MAP2 and p-cPLA2 intensity at soma and somatodendritic area. MAP2 was observed to colocalize with p-cPLA2α. The cPLA2 inhibitor ASB14780 partially suppressed these processes, suggesting that Aβ42O promotes the aggregation of MAP2 in the soma through cPLA2α activation. This finding may resemble the presence of MAP2-associated neuritic plaques found in the human AD brain (Fig.9d).

**Figure 9.**
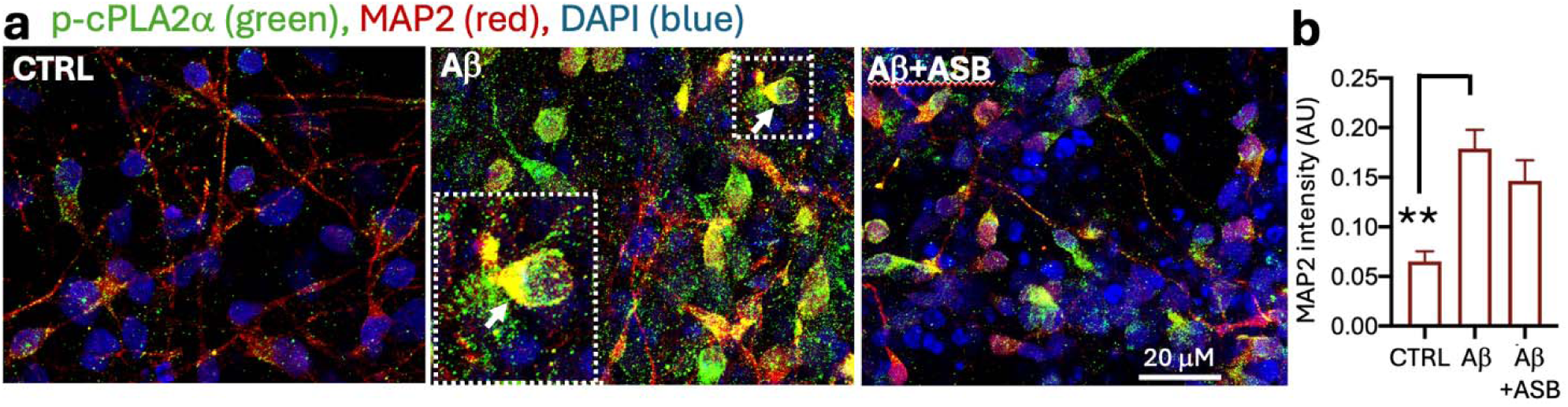
cPLA2α activation mediates Aβ42 oligomer-induced MAP2 soma aggregation with partial inhibition by cPLA2 inhibitor ASB14780 in human iPSC-derived neurons. Aβ42O increased MAP2 (red) and p-cPLA2 (green) intensity. p-cPLA2α was colocalized with MAP2 (yellow color, arrow). The enlarged box highlighted the colocalization between p-cPLA2a and MAP2. One-way ANOVA, Tukey’s multiple comparisons test. Data are presented as mean ± SEM, *\*\*p* < 0.01

### Aβ42 oligomers triggered cPLA2α activation, increased PSD-95 levels, and promoted their colocalization, which was reversed by ASB14780 in human iPSC-derived neurons

In the human iPSCs-derived neurons carrying the APOE4/4 genotype, we also investigated the impact of cPLA2 activation on postsynaptic protein PSD95. After NPCs were differentiated for four weeks, the cells were exposed to 2.5 μM Aβ42O, with or without 1 μM of the cPLA2α inhibitor ASB14780 (ASB) for 72 hours. Compared to control untreated cells, Aβ42O increased both p-cPLA2α and PSD-95 levels and promoted their colocalization in the iPSC-derived neurons (**Fig. 10**). PSD-95 was highly concentrated on the cell membrane with less dendritic distribution. These processes were partially suppressed by cPLA2α inhibitor ASB14780. In the same experiment, we also observed Aβ42O-induced reduction of CaMKIIα-labeling dendritic density and synapsin-labeling synaptic puncta and ASB treatment effects (data not shown). These data suggest that the activated cPLA2α binding to PSD-95 may mediate Aβ42O-induced excitotoxicity.

**Fig. 10.**
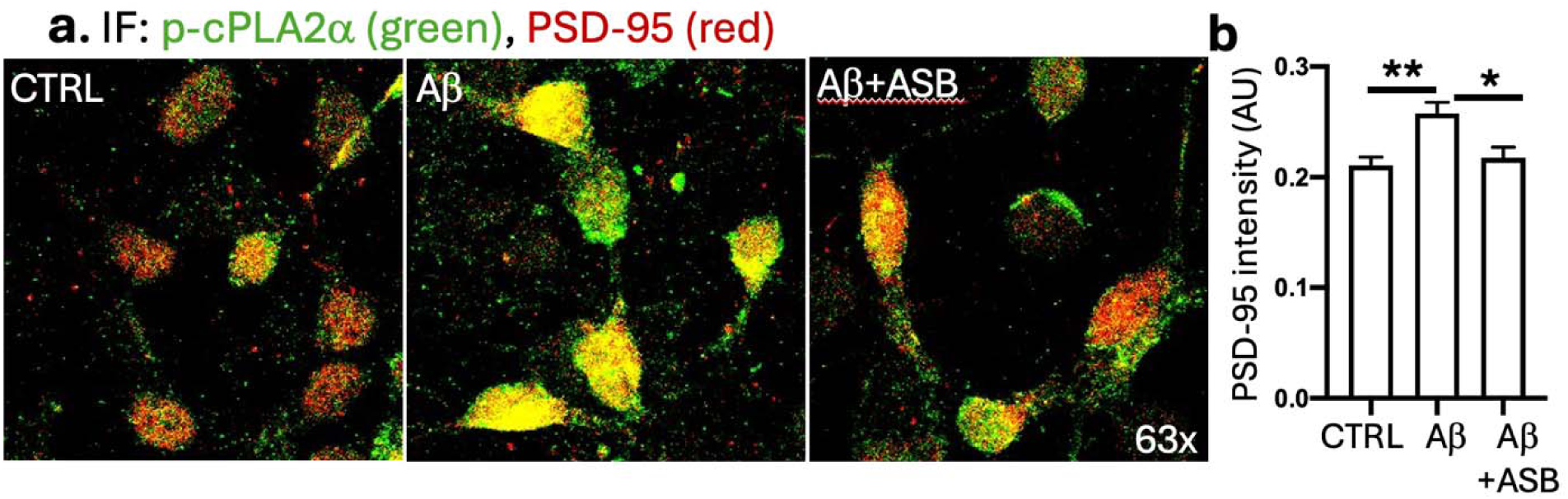
A**c**tivation **of cPLA2a induced by Ab42 oligomers increases PSD-95 and is reduced by a cPLA2a inhibitor ASB.** Aβ42 oligomers (Aβ) increased p-cPLA2α (green, **a, b**), PSD-95 (red, **a, c**) and promoted their colocalization (yellow, A) in human iPSC-derived neurons, which was suppressed by a cPLA2α inhibitor ASB. One-way ANOVA, Tukey’s multiple comparisons test. Data are presented as mean ± SEM, *\*p < 0.05, **p < 0.01*.

## Discussion

In this study, we report the first evidence of increased cytosolic phospholipase A2α (cPLA2α) and cPLA2β levels and activity in synaptosomes isolated from postmortem brain of individuals with AD, and with additional increased cPLA2β in those with MCI. The protein levels of cPLA2, including cPLA2α, phosphorylated-cPLA2α (p-cPLA2α), and cPLA2β, were negatively correlated with global cognition, and with several of the cognitive domains, including episodic memory. Additionally, p-cPLA2α was colocalized with key excitatory synaptic markers Ca^2+^/calmodulin-dependent protein kinase II subunit alpha (CaMKIIα) and dendritic marker MAP2 in cortical layer V/VI of AD and NCI. Notably, p-cPLA2α intensity was increased at synaptic clusters in AD, which was accompanied by a decrease in CaMKIIα and MAP2 intensity, suggesting a pathological shift in synaptic integrity. Moreover, both cPLA2α and cPLA2β were positively corelated with excitatory postsynaptic protein PSD95, and p-cPLA2α highly colocalized with PSD-95 in synaptosomes. In cultured human iPSC-derived neurons, Aβ42O activated cPLA2α, causing p-cPLA2 translocation to the membrane and colocalization with CaMKIIα, MAP2 and PSD95. Taken together, these data suggest that p-cPLA2α may be a key driver of synaptic dysfunction in AD pathology, and potentially of the clinical expression of AD, namely cognitive impairment.

The role of cPLA2α activation has been extensively studied in several inflammatory diseases^56^, including traumatic brain injury that causes a brain pathologies including tauopathy from chronic traumatic encephalopathy is well knows to be a risk factor for cognitive decline and dementia^57,58^. Activation of cPLA2α catalyzes the release of AA from phospholipids at sn2 acyl position^4,5^, which can subsequently be metabolized to form inflammatory eicosanoids^6,7^, such as PGD2, PGE2 and TXB2. This process is tightly regulated by Ca^2+^ concentration and site-specific phosphorylation events^20–22,59,60^. cPLA2α is mainly phosphorylated at Ser-505 by MAP kinases^13^ that play crucial roles in the development of AD pathology, such as neuroinflammation and tau hyperphosphorylation^45,61^. Consistent with these known mechanisms, we observed increased levels both cPLA2α and cPLA2β, AA and AA metabolites in AD synaptosomes. Further, high cPLA2 levels in synaptosomes correlated with low cognitive performance in ROS. This study is the first to directly link brain cPLA2 levels to cognitive function in human brain tissues, pointing to a role of cPLA2 in the clinical expression of AD.

Our data indicates that cPLA2α activation (p-cPLA2α) leads to its translocation from the synaptic cytosol to the synaptic membrane where it interacts with synaptic proteins, such as CamKIIα and PSD-95 and the dendritic membrane marker MAP2, as we demonstrated *in vivo* and *in vitro*. Localization of p-cPLA2α at the synaptic and dendritic membrane may be a key contributor to synaptic dysfunction, as CamKIIα is essential for learning, memory consolidation, and synaptic transmission^53,62^. CamKIIα dysregulation in AD is well documented^63^ and our findings of reduced CamKIIα and p-CaMKIIα in AD synaptosomes support this role. Additionally, the colocalization of p-cPLA2α with PSD-95 in synaptosomes indicates that p-cPLA2α activity is localized at excitatory postsynaptic sites, further implicating its involvement in synaptic pathology. PSD-95 is vital for synaptic maturation, stabilization and the trafficking of N-methyl-D-aspartic acid receptors (NMDARs) and α-amino-3-hydroxy-5-methyl-4-isox-azoleproprionic acid receptors (AMPARs) to the postsynaptic membrane^64,65^. PSD-95 was also increased in AD brain ^66,67^, CSF^68^, and in synaptosomes as demonstrated here, suggesting that p-cPLA2 may mediate excitotoxicity in AD^69^. Thus, p-cPLA2α may disrupt PSD-95 function, impair NMDARs and AMPARs trafficking and contribute to synaptic transmission deficits **(Fig. 11),** which are fundamental for learning and memory processes.

**Fig. 11.**
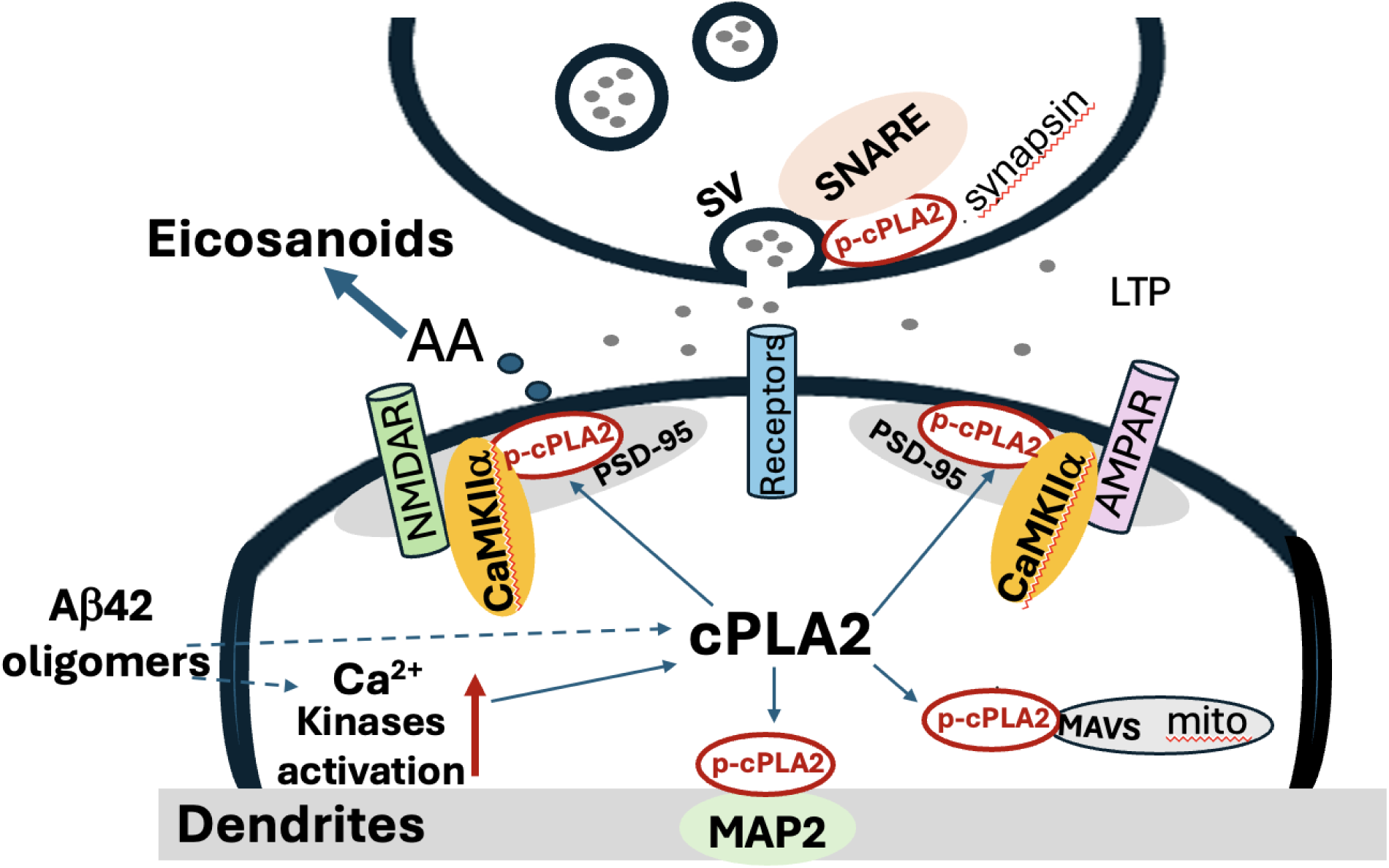
Proposed mechanism for cPLA2 overactivation causing synaptic dysfunction in AD. When activated, p-cPLA2 moves to the synaptic membrane, where it attaches to phospholipids near PSD-95 and CaMKIIa. At this site, it catalyzes the breakdown of phospholipids, producing AA. This process undermines the structural integrity and protein functionality of the synaptic membrane, affecting CaMKIIa-NMDAR and AMPAR signaling pathways during LTP. cPLA2 activation may also cause damage to presynaptic and synaptic vesicles (SV) or mitochondrial membranes, disrupting their normal functions.

The pathogenesis of sporadic AD involves complex interactions of multiple risk factors, including several which can activate cPLA2α, notably inflammatory mediators, elevated plasma homocysteine^70^, and Aβ42O. Indeed, there is considerable evidence that cPLA2α mediates Aβ oligomer-induced MAP kinase activation (ERK, JNK, P38)^71,72;32,73^, mitochondrial dysfunction^74^, tau phosphorylation^29,75,76^, synaptic loss^30,77^, neuronal apoptosis^78^, as well as CDK5/p25-induced neuroinflammation ^8,15^ in cellular and animal models. Additionally, inhibition of cPLA2α has been shown to attenuate cPLA2α-mediated-pathologies^30,31^. cPLA^-/-^ mice are resistant to soluble Aβ42 oligomer-induced synaptic and cognitive impairments^30^. Consistent with these studies, our study is the first to show that p-cPLA2α mediates Aβ42O-induced synaptic dysfunction in human and specifically in the context of AD. These observations suggest the potential of cPLA2 as a disease-modifying therapeutic target to mitigate synapse loss and cognitive impairment in AD. While amyloid-targeting therapies have shown some promise, they have been met with limitations in efficacy. In contrast, cPLA2α inhibition may provide an alternative or complementary strategy by directly addressing synaptic dysfunction and neuroinflammation, two central features of AD pathology. Moreover, because cPLA2α plays a critical role in the production of inflammatory mediators and in the regulation of synaptic protein, its modulation could potentially improve cognitive outcomes by reducing neuroinflammation and also preserving synaptic integrity. Given the complex and multifactorial nature of AD, combining cPLA2α inhibitors with other disease-modifying strategies, such as Aβ- or tau-targeting therapies, might lead to a more comprehensive approach with greater efficacy in slowing disease progression. However, challenges remain in translating cPLA2α inhibition to clinical settings. For example, further preclinical and clinical studies are necessary to identify the most effective therapeutic window and mitigate potential off-target effects among the many other considerations that will need to be addressed, such as the blood-brain barrier (BBB) permeability of the inhibitors. Nonetheless, these findings provide preliminary data for the rationale for the development of cPLA2α inhibitors as potential therapeutics for AD. Given that synaptic dysfunction is thought to occur early in AD pathogenesis, therapies aimed at modulating cPLA2α activity could potentially offer a means of preserving cognitive function before significant neuronal loss occurs. Indeed, targeting cPLA2α during early disease stages could improve the chance of delaying (or maybe even preventing the onset of) cognitive impairment in AD. The correlation of cPLA2 with episodic memory further supports this.

While our study provides evidence supporting the role of cPLA2α in synaptic dysfunction in AD, several limitations need to be addressed in future research. These include examining the relation of cPLA2 with change in cognition over the years, testing for the impact of *APOE* genotype, exploring for sex differences, and evaluating the effect of various disease stages on cPLA2α - all in a larger sample size.

## Conclusion

In conclusion, our study reveals cPLA2α activation as a key driver of synaptic dysfunction and is associated with cognitive function in AD. By associating cPLA2α with key postsynaptic proteins such as CaMKIIα, MAP2, and PSD-95, we implicate it as a potential target for synaptic loss. Inhibiting cPLA2α may be evaluated as a strategy for helping preserve synaptic integrity and slowing cognitive decline, particularly in the early stages of AD.

## Abbreviations

AD: Alzheimer’s disease
cPLA2: calcium-dependent phospholipase A2 (cPLA2α and cPLA2β)
ROS: Religious Orders Study
NCI: no cognitive impairment
MCI: mild cognitive impairment
Aβ42O: amyloid-β 42 oligomers
PSD-95: postsynaptic density protein 95
p-cPLA2α: phosphorylated cPLA2α
CaMKIIα: Ca^2+^/calmodulin-dependent protein kinase IIα
MAP2: dendritic microtubule-associated protein 2
AA: arachidonic acid
iPSCs: Induced pluripotent stem cells
PMI: postmortem interval
USC: the University of Southern California
TEM: transmission electron microscope
LDH: lactate dehydrogenase activity
WB: Western blotting
NPCs: Neural progenitor cells
KO: knockout
NMDARs: N-methyl-D-aspartic acid receptors
AMPARs: α-amino-3-hydroxy-5-methyl-4-isox-azoleproprionic acid receptors
SVs: synaptic vesicles
Syn: synaptosomes
ROSMAP_: Religious order study/memory aging project
n-RNA seq: Single-nucleus RNA sequencing
NFT _: Neurofibrillary tangles
GFAP _: Glia fibrillar acid protein
Iba1 _: Ionized calcium-binding adapter molecule 1
LC-MS/MS: Liquid chromatography–mass spectrometry
A.U. _: Arbitrary unit

## Author contributions

HNY and QLM contributed to the conception and design of the experiments. QLM performed most of the experiments and wrote the draft of the manuscript. BE, DD and SL performed lipidomic assay and reviewed the manuscript. BL analyzed lipidomic data and the single-nucleus RNA sequencing data. AS performed synaptosome isolation and BCA protein assay. JL and SW performed West Blotting. JG and MR performed immunofluorescent staining of synaptosomes and brain sections. BEK cultured mixed human iPSC-derived neurons and astrocytes. PS imaged synaptosomes under TEM. DH, AEH, DB and ZA provided brain samples and reviewed the manuscript. BEH, KMH, BGG, AL, and HCC reviewed the manuscript.

## Acknowledgements

We thank Terri L Stephen for help with editing the manuscript.

## Funding

This work was support by grants from R01AG082362 (HNY, ZA), P30AG066530 to HNY; P30AG066530 subaward (QLM); R21AG089611(QLM) P30AG066530 to Neuropathology Core, R01AG070255 (AL) ROS is supported by P30AG10161, P30AG72975, and R01AG15819.

ROS resources can be requested at https://www.radc.rush.edu and www.synpase.org.

## Availability of data and materials

All data are available in the main text or supplementary materials. Analytical codes are available from the corresponding author upon request. Sharing data from ROSMAP will follow ROSMAP data-sharing policies.

## Declarations

### Consent for publication

All authors read and approved the final manuscript.

