## Supplementary Material for "Evidence for cPLA2 activation in Alzheimer’s Disease Synaptic Pathology"

**Validation of the isolated synaptosome-enriched fraction from frozen postmortem human brain tissue**

To validate the specificity and integrity of our protocol to isolate synaptosomes, we performed Western immunoblotting (WB), LDH assay, IF staining, and transmission electron microscopy (TEM). WB analysis revealed higher levels of anti-PSD-95 antibody immunoreactivity in the synaptosome-enriched P2 fraction, but not in gliosome-enriched P1 fraction (**Fig. S1a)**. In contrast, anti-astrocyte marker GFAP and microglia marker IBA1 antibodies showed higher intensities in the P1 fraction. Synaptosomes integrity was confirmed by LDH assay (**Fig. S1b**), immunofluorescent staining using anti-PSD-95 antibody (**Fig. S1c**), and TEM (**Fig. S1d**). TEM also showed the loss of synaptic vesicles (SVs) within synaptosomes (Syn) in an 80-year-old female with AD (PMI 3.8 hours) compared to a 95-year-NCI female (PMI 3.2 hours, **Fig. S1d**).

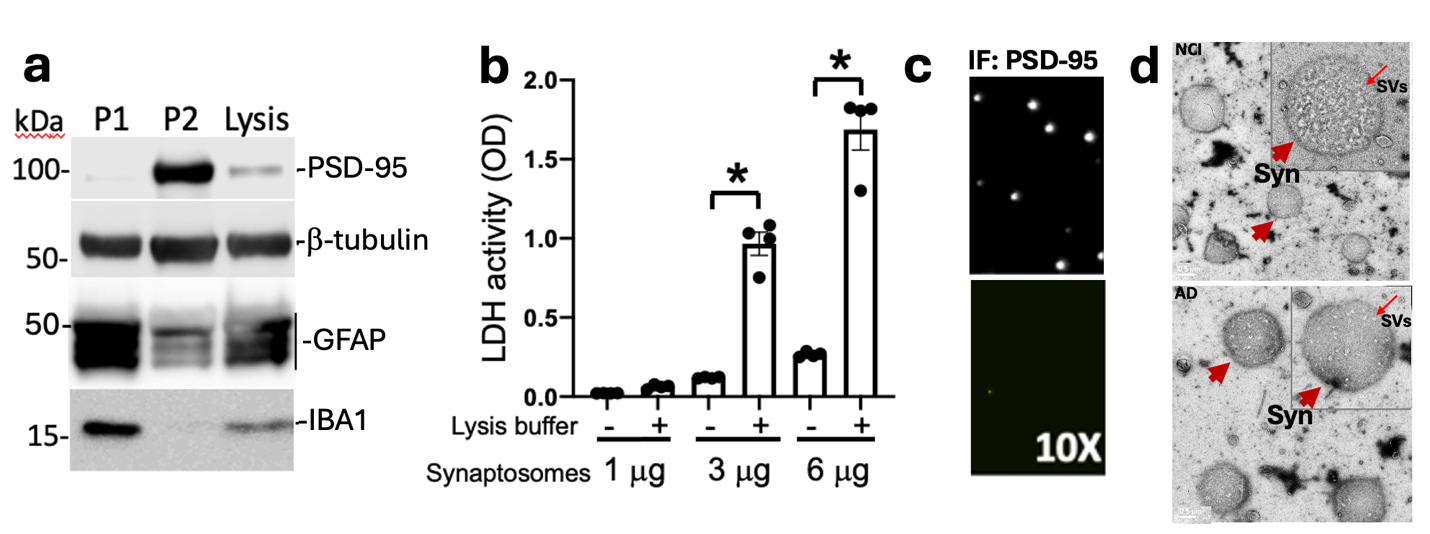

**Fig. S1 Characterization of synaptosome preparation (synaptosome-enriched P2 fraction) from frozen postmortem midfrontal cortical brain tissue. a.** Western blot **analysis** shows enrichment of the postsynaptic protein PSD-95 in the synaptosome fraction (P2) compared to the gliosome fraction (P1) and whole brain lysate. In contrast, the astrocyte marker GFAP and microglial marker Iba1 are elevated in the gliosome fraction (P1) compared to P2 and brain lysates, demonstrating successful separation of synaptic and glial components **b.** Lactate dehydrogenase (LDH) assay confirms the structural integrity of the isolated synaptosomes in the P2 fraction, indicating they remain intact post-isolation. **c.** Immunofluorescence (IF) staining of PSD-95 in synaptosomes, visualized by fluorescence microscopy, confirms the enrichment of synaptic structures in the P2 fraction. **d.** Transmission electron microscopy (TEM) image of an intact synaptosome from an 80-year-old female AD patient (PMI 3.8 hours) shows the presence and reduction of synaptic vesicles (SVs), further validating the successful isolation and preservation of synaptic elements within the P2 fraction, compared to a 95-year-old no-cognitive impairment (NCI) female donor (PMI 3.2 hours).

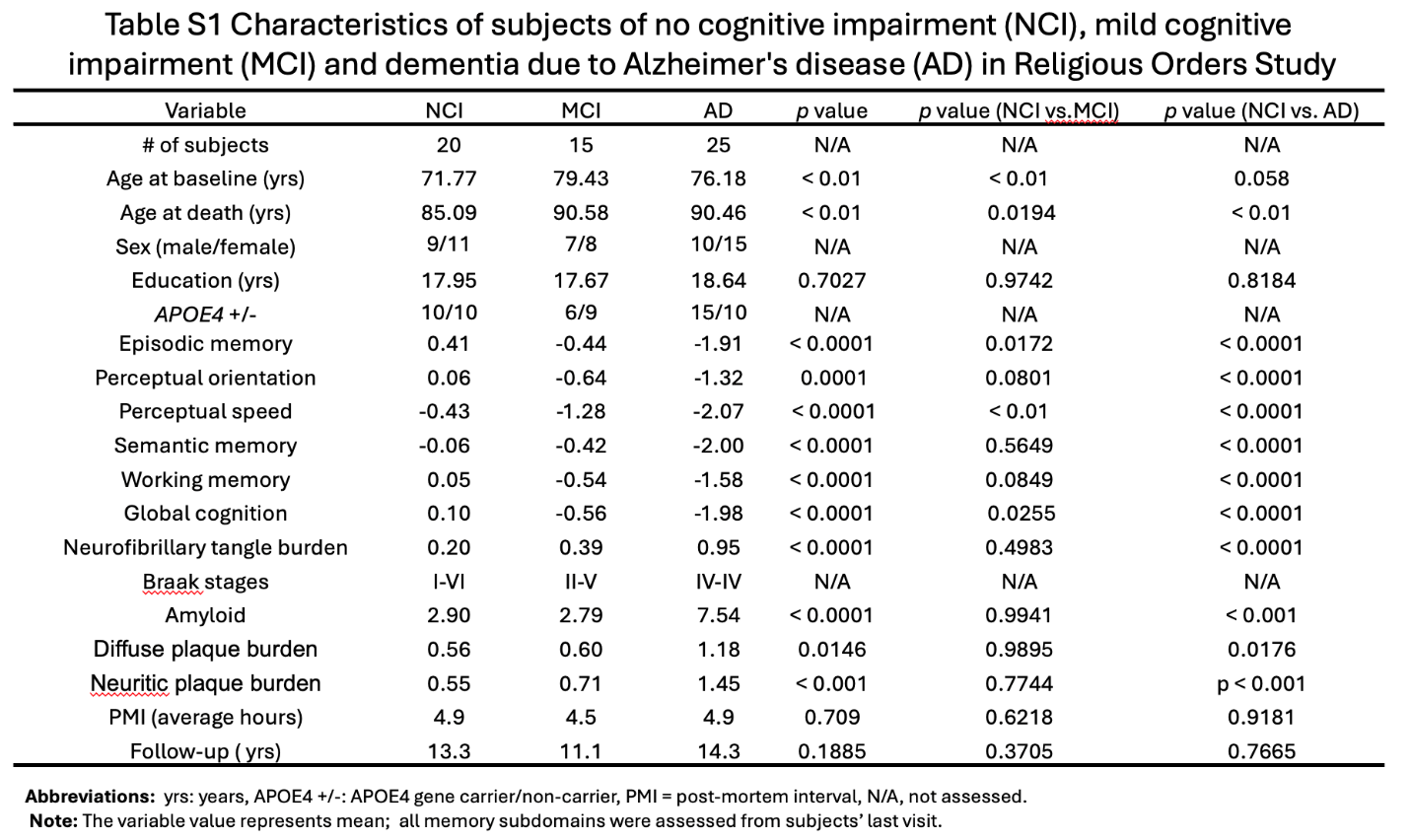

Abbreviations:  yrs: years, APOE4 +/-: APOE4 gene carrier/non-carrier, PMI = post-mortem interval, N/A, not assessed. Note: The variable value represents mean; all memory subdomains were assessed from subjects’ last visit.

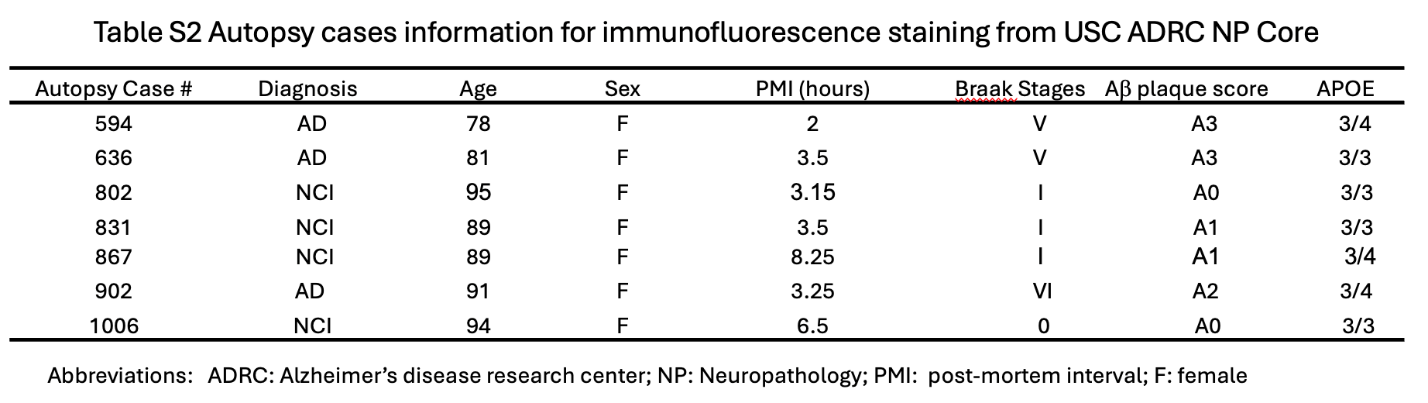

Abbreviations:   ADRC: Alzheimer’s disease research center; NP: Neuropathology; PMI: post-mortem interval; F: female.

Abbreviations:   WB: West blotting, IF: Immunofluorescent staining 
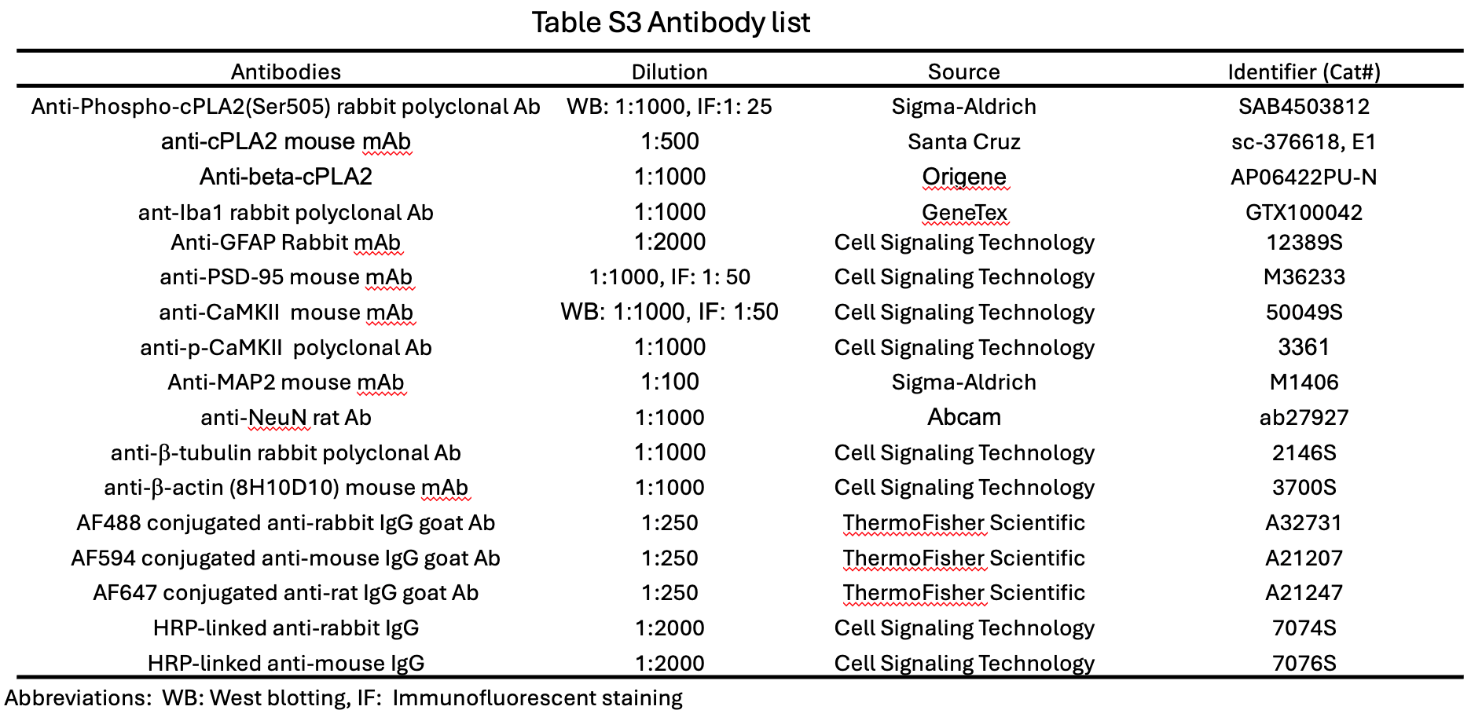
